# Rapid Response of Neutrophils Determined by Shortest Differentiation Trajectory and New Insight into Cell Cycle Kinetics and Chronological Order of Progenitor States

**DOI:** 10.1101/2024.04.23.590178

**Authors:** Yongjian Yue

**Author notes:** **Correspondence:** Yongjian Yue Address: Shenzhen People’s Hospital, No. 1017 Dongmen North Road, Luohu District, Shenzhen 518020, China.

## Abstract

Despite substantial advances in our understanding of hematopoietic stem cell self-renewal, differentiation, and proliferation, the underlying mechanisms and cell cycle kinetics remain elusive. Neutrophils, as primary rapid responders during infections, quickly generate an enormous number of mature cells for immune defense. This immune phenomena is effected by cell cycle kinetics. Our previous study redefined progenitors and reformed the hierarchy of hematopoiesis. How exactly lineage commitment and cell cycle are coordinated remain to be elucidated. Here, we aim to elucidate the cell cycle kinetics and chronological order of progenitor states, providing new insights into immune phenomena. We revealed the changes in cell cycle kinetics from differentiation (lineage commitment) to proliferation (cell division) in peripheral hematopoiesis. Differentiating hematopoietic progenitor cells (HPCs) are maintained in the G1 phase of cell cycle, regulated by DNA replication. CDCA7 was identified as an essential factor for DNA replication, facilitating the transition of progenitor cells from G1 to S phase. Once the progenitors complete their commitment, cell cycle states of progenitors convert from G1 to S phase, switching from lineage commitment to cell division with DNA replication begins. Lineage commitment and cell division of MPCs are independent processes, and fate determination takes precedence over proliferation. Since the differentiating HPCs do not undergo cell division, a committing progenitor will destined to mature into only one committed progenitor cell along its designated differentiation trajectory. The chronological order of quiescence, self-renewal, differentiation, and proliferation, was clearly delineated. Our study proposed the concept of "**temporal sequence control**," emphasizing that fate determination necessarily precedes division during lineage commitment, thereby providing a new dimension for modeling hematopoietic dynamics. Our study reveals that initial progenitors in adult peripheral blood exhibit self-renewal capabilities, which are absent after lineage commitment is completed during hematopoiesis. Neutrophil present the shortest differentiation trajectory in the hematopoietic hierarchy, which decide their role as rapid responders. Our results provide a clear characterization of the cell cycle kinetics of HPCs’ lineage commitment and cell division, facilitating advancements in inflammation and disease treatment.

Graphical Abstract:
Lineage commitment progresses independently of cell division, where each progenitor differentiates into a single terminally committed cell (Panel B), **challenging** the conventional paradigm that cell division occurs at each step of lineage commitment (A).s

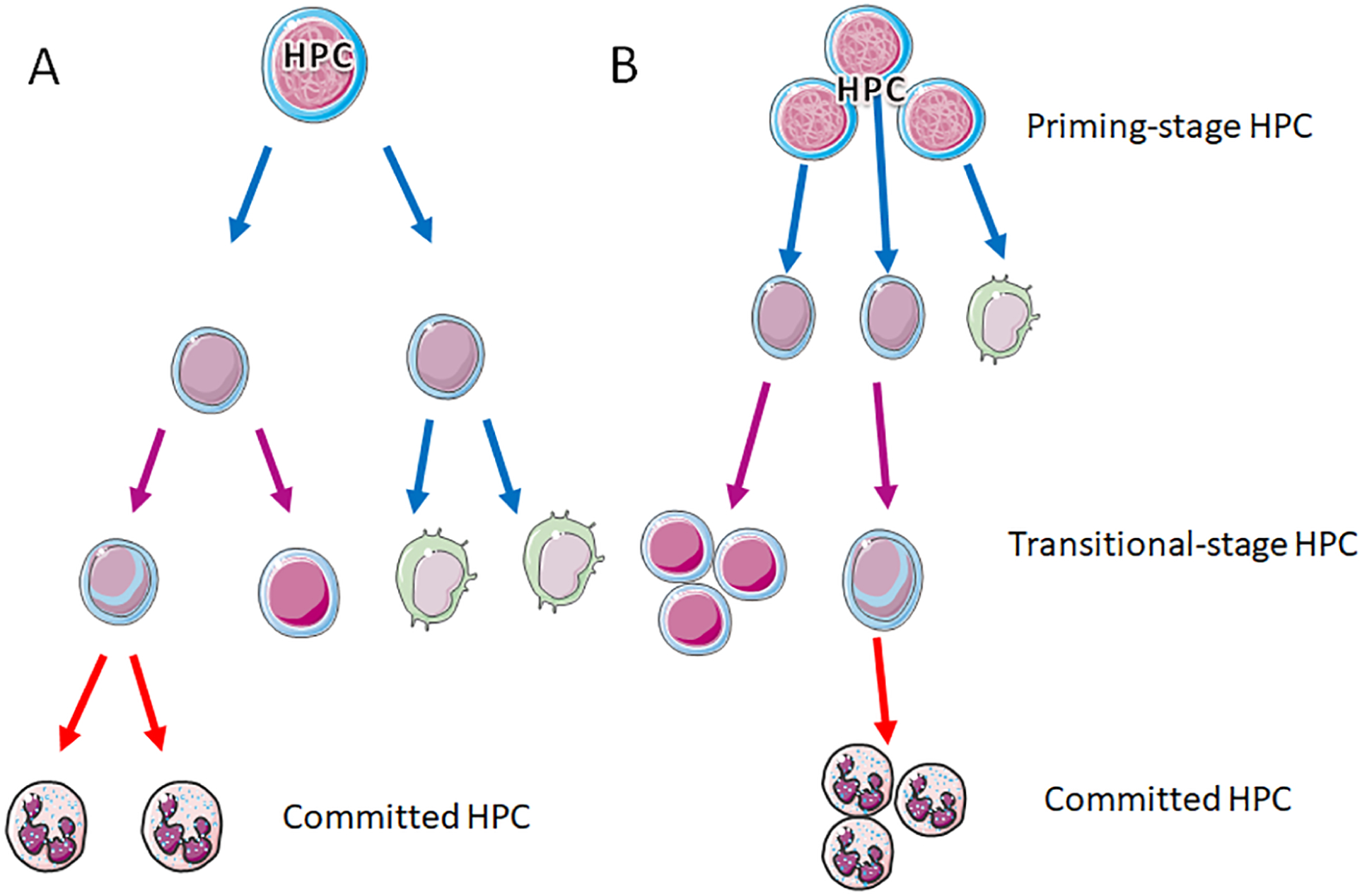

## Introduction

Hematopoietic stem cells (HSCs) possess the potential of self-renewal, generating daughter HSCs by cell division, and differentiating into various blood cell lineages. Despite substantial advances in our understanding of HSC self-renewal and differentiation, the cell cycle kinetics and underlying mechanism remain elusive[1]. Inefficiently capture and lineage heterogeneity impede standardized definitions and the revelation of molecular mechanisms underlying fate decision and lineage commitment [2, 3]. Moreover, single-cell RNA (scRNA) sequencing transcription atlas attempts to redefine progenitors and reform the classical hematopoietic hierarchy, but contradictions persist [4, 5]. These contradictions are influenced by progenitor original, capture methods, inefficient definitions, and analysis algorithms [6]. It widely accepted that progenitors continuously differentiate into all lineages during hematopoiesis [7, 8]. This continuous differentiation makes it difficult to define transitioned progenitors by surface marker due to the lack of reliable boundaries among them [5]. Therefore, owing to inconsistent definitions and conflicting hematopoietic hierarchical models [9], the cell cycle kinetics and regulatory mechanisms underlying self-renewal and lineage commitment remain unclear. Investigating the state transition from quiescent to activation during hematopoiesis remains challenging.

Our previous study redefined the entire spectrum of original hematopoietic progenitor cells (HPCs) and reformed the hematopoietic hierarchy roadmap using a comprehensive transcriptional atlas [10]. Hematopoietic system can be reconstituted by HSC transplantation, suggests that the self-renewal potential exist in HPCs within peripheral blood following G-CSF mobilization. The states of differentiation, proliferation, and self-renewal of progenitors can be captured simultaneously, as progenitors continuously differentiate into all lineages during hematopoiesis. Previously study demonstrated that hematopoietic stem cells can differentiate into restricted myeloid progenitors before cell division in mice[11]. Nevertheless, the precise coordination between lineage commitment programs and cell cycle regulation remains incompletely characterized. Here, based on the clearly redefinition and reformed hierarchy of hematopoiesis, we aim to elucidate the cell cycle kinetics and underlying mechanism of self-renewal, differentiation, and proliferation during peripheral hematopoiesis. Furthermore, we purpose to provide a new interpretation of immune phenomena, such as the causation of neutrophils as the primary and rapid responders during infections. Our study showed the initial progenitors in adult peripheral blood exhibit the self-renewal capabilities but gradually diminish after lineage commitment is completed during the hematopoiesis. Progenitor should complete commitment before switched from differentiation to proliferation, a cell cycle status conversion process controlled by the DNA replication. This study makes significant breakthroughs in the fundamental discipline of immunity development. Collectively, our results highlight numerous possible avenues for achieving precise control of progenitor cell differentiation, facilitating advancements in drug development and leukemia treatment.

## Results

### The co-segregation of neutrophils progenitor cells (NPCs) was decide by GATA2 and CSF3R

Based on the efficient capture and unsupervised dimensionality-reduction analysis, all stages of HPC were redefinition by the scRNA transcriptional atlas in our previous study (Figure 1A)[10]. To delineate the two stages of neutrophilic progenitors, a re-analysis was conducted to identify marker genes for distinguishing them. We identify new efficient marker genes of neutrophilic progenitors, such as *MPO, MGST1, IGLL1, C1QTNF4*, and *NPW* (Figure 1B). The specific genes of *ELANE* and *AZU1* were rarely expressed in these stages, indicating that some markers may only be active during the mature differentiation stages, but not in progenitor stages. The initial differentiation stages of hematopoiesis simultaneously primed into megakaryocytic–erythroid, lymphoid, and neutrophilic linegaes. The characterization of *CSF3R* and *GATA2* activation clearly demonstrated the co-segregation of neutrophils progenitor cells (NPCs) with other lineages (Figure 1C). *CSF3R* expression was continuously high during the differentiation of neutrophils, but gradually restrained in the megakaryocytic–erythroid and lymphoid lineages differentiation process. *GATA2* activation was gradually up-regulated in the megakaryocytic–erythroid lineages but was suppressed in the other two lineages. The modulation of these two genes displayed a synergistic function in fate decision during the initial stage of hematopoiesis.

**Figure 1.**
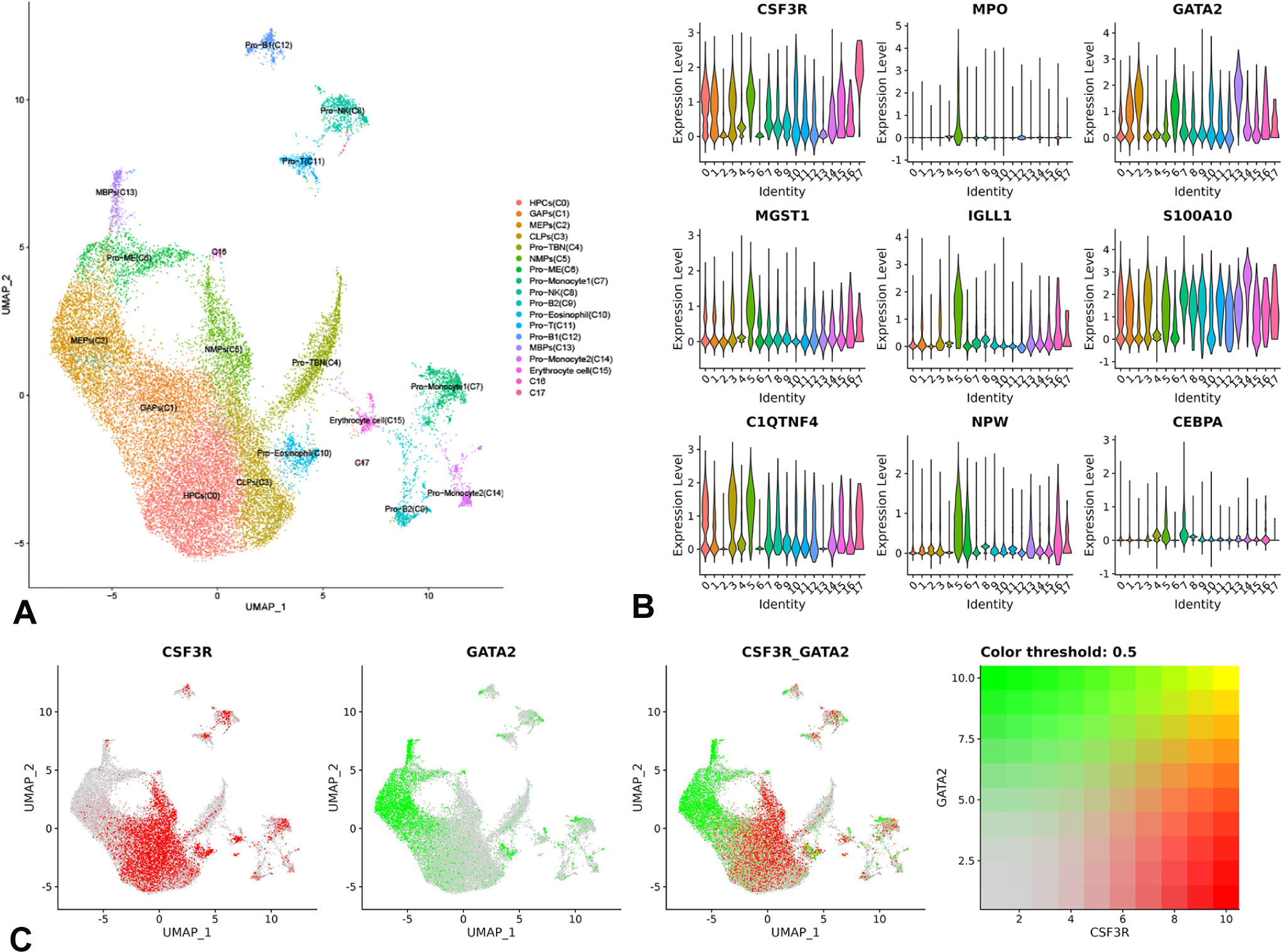
Co-segregation of neutrophils and megakaryocytic–erythroid progenitor cells commitment. A, UMAP illustrating 17 redefined clusters of adult peripheral blood CD34+ progenitor cells[10]. B, Violin plots showing the expression of neutrophil marker genes, revealling that *MPO*, *IGLL1*, and *MGST1* are more efficient for definition. C, Expression-feature plots of CSF3R and GATA2 clearly demonstrated co-segregation among neutrophils and megakaryocytic–erythroid progenitor cells. UMAP, uniform manifold approximation and projection; MPC, multi-potent progenitor cells; GAPs, gene-controlled progenitors; CLPs, common lymphoid progenitors; MEPs, megakaryocytic–erythroid progenitors; MBPs, mast cell, basophil, or eosinophil progenitors; Pro-MEs, megakaryocytic and erythroid progenitors; NMPs, neutrophilic and monocytic progenitors.

### The shortest differentiation trajectory and minimal signaling changes of NMPs commitment

Cell cycle analysis was conducted to confirm the transformation during hematopoiesis. Our results showed that most progenitors of NMPs, MBP, and Pro-ME were already in the S phase, while lymphoid progenitors were not (Figure 2A). This indicates that neutrophil progenitors are almost complete commitment and shift to proliferation after initial priming from multi-potent progenitor cells (MPCs). In contrast, megakaryocytic–erythroid progenitors undergo two transitional stages (GAP and MEP) before transitioning to the proliferation stage (Pro-ME). Compared to the MEP lineage, differentiation process of neutrophils is the shortest in hematopoiesis (Figure 1A). The expression of the MCM complex genes, component of the pre-replication complexes (pre-RC), was highly expressed in the late stages of the megakaryocytic–erythroid (Pro-ME), mast cell-basophil progenitors (MBPs), and neutrophilic lineages (NMPs) (Figure 2B). This confirmed that these three lineage progenitors switched from differentiation (lineages commitment) to proliferation (reproduction). The lineage commitment of these three progenitor types is a continuous process without distinct boundaries. Furtheremore, the proportion and number of neutrophil progenitors rank among the top three lineages, explaning why neutrophils are the second most abundant cell type in whole blood, only behind to erythrocytes. The expression characterization of CSF3R, highly expressed throughout the differentiation trajectory of NMPs (Figure 1C), indicates that CSF3R-activated signaling may be highly consistent with neutrophils differentiation signaling. Furthermore, the activation of unique key TFs is required for the lineages commitment of MEP with *GATA2* and CLP with *HOPX*. However, no additional distinct TFs are identified as necessary for NMPs commitment. This suggests that the transformation of initial progenitors (MPCs) into neutrophilc progenitors occurs rapidly with minimal and low-threshold signaling changes. The shortest differentiation trajectory and minimal signaling changes ensure a quick response and the rapid generation of an enormous of neutrophils.

**Figure 2.**
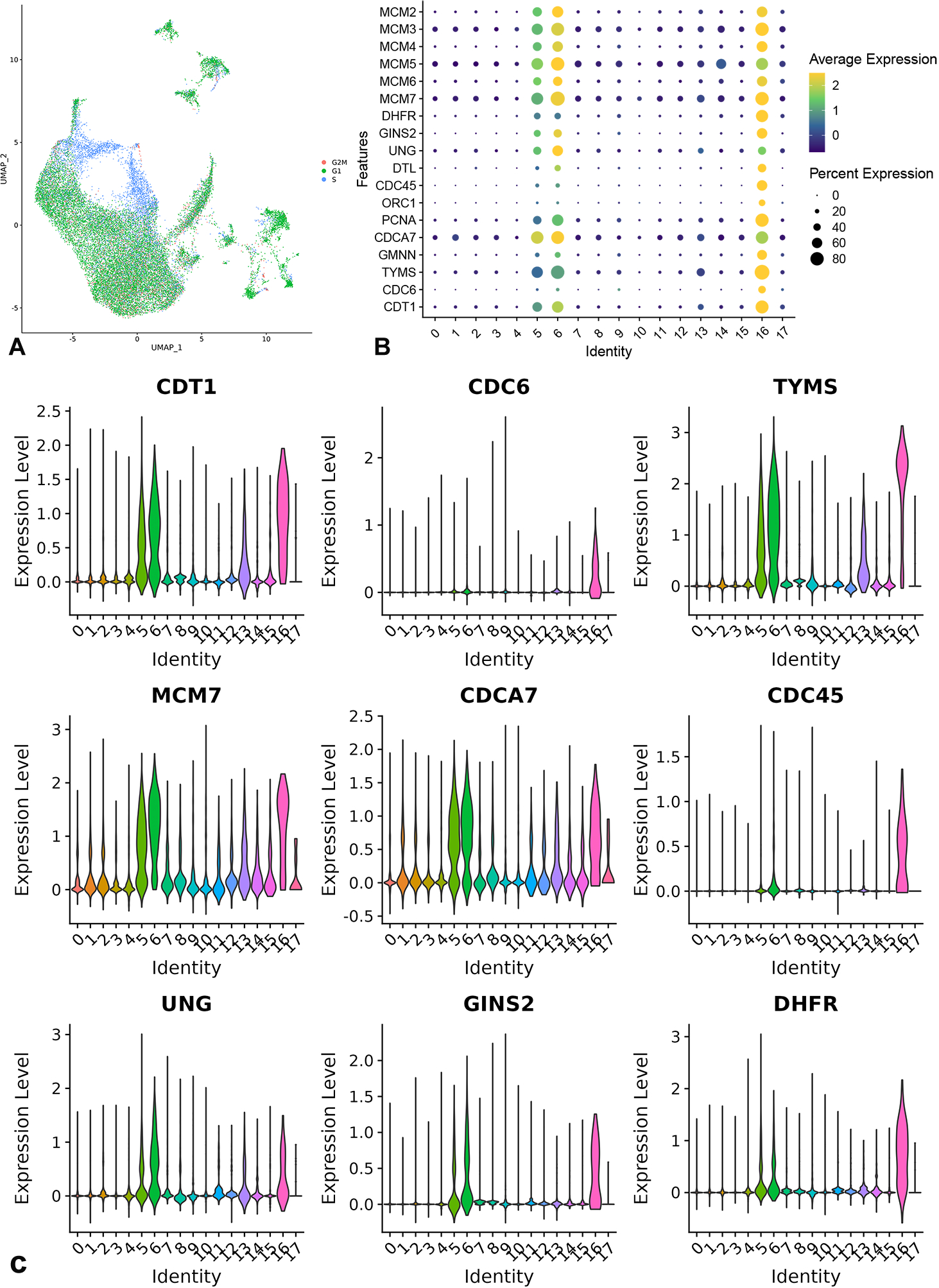
Cell cycle analysis and characterization of cell cycle markers genes. A, UMAP illustrating the cell cycle phases of progenitor cells, displaying that the NMPs, MBPs, and Pro-ME are already in S phases in adult peripheral blood. B, Dot plots showing the expression of representative cell cycle markers genes. C, Violin plots displaying the expression of cell cycle markers and unique marker genes, indicating that *PCNA*, *TYMS*, and *CDT1* were up-regulated in S phase progenitors.

### The CDCA7 play a key role during the genes determining cell cycle kinetics transformation from commitment to proliferation of progenitors

Since the switch from G1 to S phase in progenitors was identified, we aimed to identify the genes that control cell cycle progression and differentiation. Quiescent hematopoietic progenitors were maintained in the G0 phase of the cell cycle. G-SCF mobilization triggered the switch from a quiescent to an activated state, indicating lineages commitment or self-renewal activity. The genes of *CDT1, TYMS, UNG, DHFR, PCNA, GINS2*, and *DTL* (CDT2) were highly expressed in the S phase progenitors (Figure 2B)[12]. Interestingly, the cell division cycle gene *CDCA7* was significantly up-regulated during the progenitor cell cycle. However, the genes *CDC6* and *CDC45*, which encoding proteins essential for the initiation of DNA replication, were rarely expressed in S phase progenitors compared with *CDCA7*(Figure 2C). This indicates that *CDCA7* may play a critical role in the initiation of DNA replication in hematopoiesis, unlike CDC7[13]. Furthermore, the up-regulation of these cell cycle genes confirmed that most NMPs and Pro-ME were already in the S phase, indicating a switch from differentiation to proliferation (Figure 2A, B). By sequentially expressing DNA replication initiation elements, the transitioned progenitor cells were primed into final differentiation stages and completed lineage commitment (Figure 2A). CDK4 and CDK6 were highly expressed in all progenitor lineages, and CDK4 being up-regulated in S phase of progenitors (Figure 2B). However, CDK1 was absent in all progenitors, and CDK2 was uniquely expressed in Pro-ME. The Pro-ME stage had already initiated the CDT1 destruction process of S phase, characterized by high expression of PCNA, GMNN, and DTL (CDT2, CRL4) (Figure 2B). The CRL4 (CDT2) complex mediates the polyubiquitination and subsequent degradation of CDT1 and other cell cycle regulated genes. The transformation of uracil to thymine was very important during DNA replication. Our results showed that three enzymes involved in thymidylate synthase were up-regulated, including *DUT, TYMS*, and *DHFR* (Figure 2B). DUT catalyzes the cleavage of dUTP to dUMP, and TYMS catalyzes the reductive methylation of dUMP to dTMP. Uracil DNA Glycosylase (UNG) plays an important role in preventing mutagenesis by eliminating uracil from DNA molecules. In summary, the expression of DNA replication-controlled genes of the cell cycle indicated that cell proliferation predominantly occurred in NMPs, MBP, and Pro-ME, but rarely present in other progenitor clusters.

### Absence of progenitor self-renewal following lineages commitment in adult peripheral blood

HSCs were maintained in the G0 phage, remaining quiescent state in the bone marrow. The mechanism that drives progenitor cells into cell cycle remains unknown, since the challenge of distinguishing the molecular activities involved in commitment, self-renewal, and proliferation. Our results showed that the majority of the progenitors in the three clusters (NMPs, Pro-ME, and MBPs) had already entered the S phase of cell cycle, which was markedly different from other initial and transitional state clusters (Figure 2A). Most progenitors in the initial and transitional state clusters were maintained in the G1 phase of activated state, indicating that these progenitors tend to undergo lineages commitment without cell division (Figure 2A). **It is impossible for progenitors lacking DNA replication activity to have self-renewal ability**. Conversely, the progenitors in the S and G2/M phases in the initial and transitional state clusters may engage in asymmetric or symmetric self-renewal division, maintaining their self-renewal potential. Approximately 30% of the rare cells in these clusters that entered into S and G2/M phases exhibited self-renewal division activity. The expression of cell cycle genes was highly activated in most NMPs, Pro-ME, and MBPs (Figure 2B). This reminds us that during the lineage commitment of progenitor cells, each lineage progenitors should complete the differentiation (lineages commitment) process before transitioning to proliferation (cell division). Further Gene Ontology (GO) and Kyoto Encyclopedia of Genes and Genomes (KEGG) signaling results showed that DNA replication initiation and cell cycle DNA replication were significantly enriched in NMPs, MBPs, and Pro-ME (Figure 3A, B). Additionally, we identified only low signaling of DNA replication activation in the original progenitor group of MPCs. Progenitors that even do not exhibit signaling of DNA replication activity do not possess self-renewal ability. The results revealed that along the trajectory of hematopoiesis, progenitors gradually differentiate into committed stages and transition into DNA replication of cell cycle before proliferation (Figure 3C). DNA replication may be arrested by the activated signaling of commitment. Cell cycle kinetics regulations ensure accurate commitment process. In conclusion, HPCs were maintained in the G1 phase by controlling DNA replication initiation and the DNA replication of cell cycle. Undifferentiated progenitors maintained in the G1 phage tend to activate lineages commitment procedures, whereas those that undergo DNA replication of cell cycle exhibit self-renewal activity. In contrast, differentiated progenitors do not have self-renewal ability and transition into proliferation.

**Figure 3.**
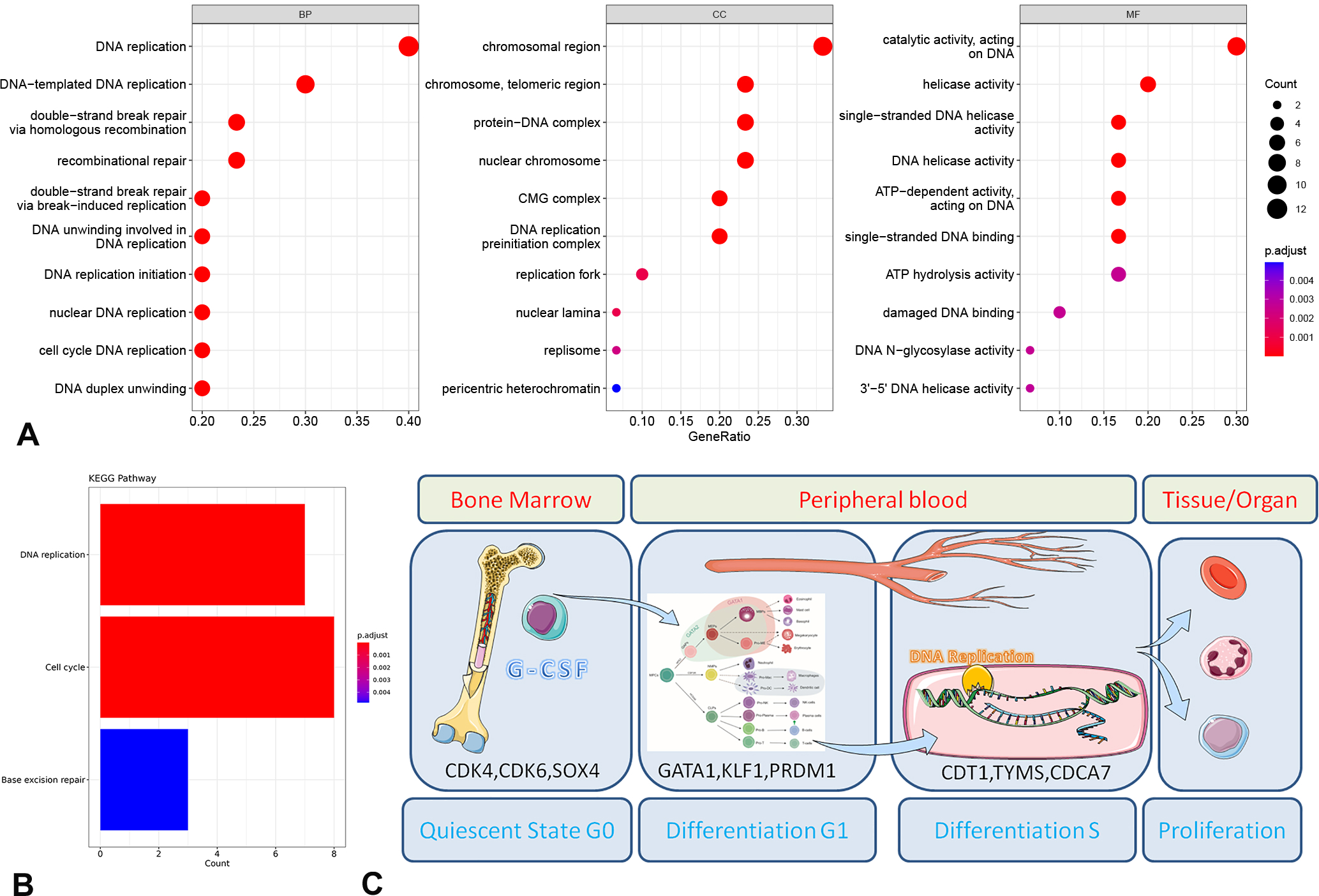
GO and KEGG analysis of cell differentiation. A, Bar plot displaying enriched categories in the GO analysis, showing significant enrichment in DNA replication initiation and DNA replication of the cell cycle. B, Enrichment results of three signaling pathways in progenitor cells. C, Schematic showing the switch of progenitor cell from a quiescent state to differentiation and proliferation of the cell cycle, illustrating that differentiation occurs prior to proliferation.

### Governs the cell cycle kinetics of activated progenitor cells: self-renewal, lineages commitment, proliferation, or re-entering the quiescent state

Activated progenitor cells must decide whether to enter the cell cycle for self-renewal, whether to enter differentiate of lineages commitment or whether to re-enter the quiescent state. As previously mentioned, initial activated MPCs (mobilized) are stay at G1 phase of the cell cycle, while DNA replication and proliferation mainly occurred after lineages commitment is completed in committed progenitors (Figure 3C). Lineages commitment and cell division of MPCs are independent processes. Lineages commitment of differentiation in HPCs precedes cell division of proliferation during hematopoiesis (Figure 3C). This indicates that the activated initial G1 phase progenitor cells (MPCs) is the key decision point. The cell cycle states maintaining in the G1 phase are coupled to the fate determination of progenitor cells during lineage commitment. The cell cycle states may not directly affect the fate determination of MPCs but only provide a stable activated environment for cell differentiation, since different progenitor lineages presented consistent cell cycle states conversion. Fate determination of MPCs was governed by strictly orchestrated extrinsic signals (cytokines) and intrinsic regulators (fate decision genes). Differentiation signals of committing HPCs inhibit cell cycle states conversion, arresting them in the G1 phase. This meaning that fate determination takes precedence over proliferation. Once progenitor complete commitment, cell cycle state conversion was activated, entering into S phase where DNA replication and subsequent proliferation occur. This precise sequential activation of differentiation and proliferation ensures the accurate determination and high-efficiency generation of specific cell types during hematopoiesis.

Here, we propose an integrative model to elucidate the complex chronological order among HSC quiescence, self-renewal, and differentiation (Figure 4). Firstly, actived MPCs can easily revert to a quiescent state due to minimal signaling changes required during the incomplete cell cycle transition from the G0 to G1 phase. This indicates that G1 phase MPCs reverting to the quiescent state without exiting the entire cell cycle; they arrest in the G1 phase with only a few signaling changes. With activating signals, such as G-CSF mobilization and other environment stimuli, MPCs can become activated but can re-enter the quiescent state without triggering self-renewal or differentiation (lineage commitment). Secondly, the actived MPCs infrequently enter the cell cycle for self-renewal once the conditions for DNA replication were all set (Figure 4). A low level of MPC self-renewal maintains the HSC pool size properly and the homeostasis of the hematopoietic system. However, a probability is that MPCs self-renewal may not occur frequently because DNA replication initiation genes are very low expression in MPCs state. Thirdly, G1 phase MPCs enter lineages commitment once fate decision factors of commitment were activated by physiological or emergency stimuli, such as high secretion of G-CSF, GM-CSF or EPO (Figure 4). The genetic factors, such as fate decision TFs, cooperate with the receptor signaling activation during lineages commitment. That concludes that MPCs can differentiate into progressively restricted lineage progenitors without cell division. The absence of cell division in uncommitted progenitors **challenges the prevailing paradigm that clonal haematopoiesis** is driven by HSPC clone expansion[14, 15], suggesting more complex ontogenic mechanisms underlie this phenomenon.**. The general defined clonal hematopoiesis was more like a procedure of committed hematopoietic cell division and differentiation.** The transformation of states is triggered by extracellular signals, which synergize with intracellular molecular and genetic signaling. In summary, MPCs maintained in the G1 phase possess the most favorable state for deciding between quiescence, self-renewal, and differentiation. These decisions are influenced by extracellular signals, physiological conditions, environmental factors, and intracellular genetic and epigenetic factors. Finally, once cells complete commitment, these differentiated progenitors enter the S phase, proliferate, and mature into various hematopoietic cells. The committed hematopoietic cells (restricted lineage) start proliferation (division) and further conducted second time differentiation formed various type of same kind immune cells depends on the physiologic and environmental requirement. Thus, after HPCs complete lineage commitment and their designated differentiation trajectory, they undergo extensive cell division. Their progeny then undergo subsequent functional differentiation according to physiological demands in different microenvironments. This functional differentiation-proliferation process is distinct and independent from the lineage-specifying fate determination processes observed in HPCs. In summary, the chronological order of progenitor states, including self-renewal, differentiation, and proliferation, is revealed.

**Figure 4.**
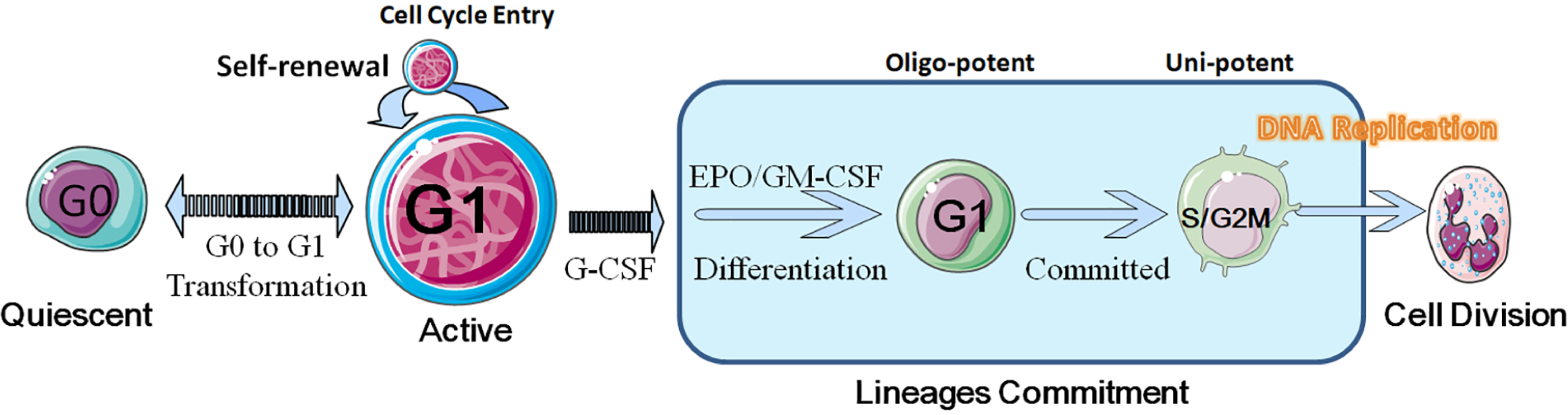
An integrative model to elucidate the cell cycle kinetics and the complex chronological order among MPC quiescence, self-renewal, and differentiation. Bone marrow HSCs remain in a quiescent state (G0). The actived MPCs can revert to quiescent state (HSC) or transition to the G1 phase with the activating of G-CSF or other environment stimuli. MPCs infrequently enter into the cell cycle of self-renewal. The G1 phase MPCs enter differentiate of lineages commitment depending on physiological or emergency circumstances, such as high secretion of GM-CSF or EPO. Once commitment is complete, these committed progenitors enter into the S phase, proliferate, and mature into various hematopoietic cells.

## Discussion

Overall, we revealed the chronological order of progenitor cell cycle state maintenance and provide new interpretations of immune phenomena through cell cycle analysis based on the clear redefinition and reformed hierarchy of hematopoiesis [10]. Our study showed that NMPs, MBPs, and Pro-ME were derived into the DNA replication phase of cell cycle, indicating they had completed differentiation of lineages commitment and transitioned to proliferation. Critical DNA replication initiation factors were identified. We further verified the shortest differentiation trajectory of neutrophils in hematopoiesis. Our findings indicted that the progenitors in adult peripheral blood, having completed lineages commitment, did not exhibit self-renewal capabilities during hematopoiesis. HPCs were maintained in the G1 phage (differentiation), controlled by the initiation of DNA replication. We revealed the cell cycle kinetics switching from differentiation to proliferation among lineage-committing progenitor cells during hematopoiesis**. Our findings revealed that lineage commitment is uncoupled from cell division, fundamentally challenging the conventional paradigm of "coupled differentiation and proliferation" and redefining the temporal hierarchy of hematopoietic regulation.**

Due to the limitation of unclear progenitor definitions, the transitional stages during hematopoiesis were unknown. Here, we detailed the transformation of progenitors from differentiation to proliferation during hematopoiesis. The mini-chromosome maintenance (MCM) complex, which includes *MCM2*, *MCM6*, *MCM5*, *MCM4*, *MCM3*, and *MCM7* genes, is a component of pre-replication complexes (pre-RC) [16]. During the G1 phase of the cell cycle, CDT1 cooperates with CDC6 and the origin recognition complex (ORC) to promote the loading of the MCM complex onto DNA to generate pre-RC [16]. The pre-RC plays an essential role in the eukaryotic cell mitosis. Our study showed the genes of pre-RC were up-regulated in the proliferation state progenitors (NMP, MBP, Pro-ME). Additionally, the GINS complex, a core component of CDC45-MCM-GINS (CMG) helicase, plays an essential role in the initiation of DNA replication and the progression of DNA replication forks [17]. Our study showed the cell cycle S phase-related genes were activated in the NMP, MBP, and Pro-ME. However, other cluster exhibited only low levels of DNA replication activity. This indicates that these three types of progenitors had transitioned from differentiation to proliferation. CDCA7 was identified as involved in the initiation of DNA replication and the progression of DNA replication, differing with CDC6 and CDC45, which initiated DNA replication. CDCA7 may be involved in determining the short-lived characterization of neutrophils.

Self-renewal of HSCs is crucial for maintaining blood system homeostasis. Generally, progenitor self-renewal must involve DNA replication during cell division, regardless of the type of division. However, our study demonstrated that most HPCs were maintained in the G1 phage (differentiation), which was arrested by the DNA replication of cell cycle. Progenitors should complete differentiation procedure (without cell division) before transitioning to the proliferation (with cell division) stage. This process is analogous to a tree that must grow to maturity before it can bloom and bear fruit. Previously study demonstrated hematopoietic stem cells can differentiate into restricted myeloid progenitors before cell division in mice[11], that was consistent with our observed phenomenon in human. Our HPCs differentiation trajectory stages and cell cycle kinetics revealed that self-renewal does not occur after lineages commitment is completed in adult peripheral blood, whereas proliferation (cell division) did. This provides solid interpretation of why progenitors cannot proliferate extensively *in vitro*. Clonal hematopoiesis can only occur in fully committed progenitor cells and their progeny, but not in hematopoietic progenitor cells (HPCs) at the commitment stage. Our conclusion challenges the classical view that ’early progenitor cells can drive clonal expansion through division.’ **Our study demonstrates that spatiotemporal constraints of clonal expansion play a pivotal role in hematopoiesis.** HSC transplantation can reconstitute a healthy blood system due to the self-renewal potential of HSC. Our previous study constructed an expression atlas utilizing both mobilized and quiescent state progenitors, yet we did not observe self-renewal after lineages commitment was completed during hematopoiesis. One possible explanation is that self-renewal occurs in uncommitted transplanted progenitors, while committed progenitor cells retain robust proliferation abilities. Another possibility is that differentiated cells (NMP, Pro-ME) may also possess self-renewal capabilities, conflicting with the notion that only early progenitors have self-renewal potential. It can be concluded that multipotent self-renewal progenitor do not exist after lineages commitment is completed in peripheral blood. Whether Pro-ME and NMPs possess self-renewal capabilities still requires further investigation. Furthermore, lineages commitment factors, which play a role in determining unique cell type fate, may not be related to self-renewal, such as SCF, GM-CSF, and TPO. Since the cell cycle occurs in differentiated lineages, this indicates that lineages-specific cytokins are not key regulators of the cell cycle.

Bone marrow hematopoietic progenitors remain in G0 phase of the cell cycle. After the initial mobilization by G-SCF, progenitors differentiated into multipotent progenitor cells, losing their self-renewal capacity [18]. Mobilized progenitors lack cell division capacity while retaining fate determination potential[19]. The mechanisms underlying the transformation from a quiescent to an activated state remain unclear. Our study showed that CSF3R, expressed in the initial subsets of progenitors, is the key signaling receptor for activation. Furthermore, our previous study revealed that the transcription factors of *SOX4, CDK4, XBP1, FOXP1*, and *CDK6* were highly expressed across all progenitor clusters during hematopoiesis [10, 20]. The expression of these TFs, which generally exhibit a synergistic relationship with CSF3R signal activation, indicates that quiescent progenitor cells were mobilized and assigned to the next stages of differentiation. We also clearly identified the lineages-specific fate decision transcription factors during lineages commitment based on the reliable redefined hematopoiesis framework [10]. Since these factors were uniquely expressed and activated in differentiation stages of progenitor but occurred later than the mobilization, this confirmed that lineages-specific fate decision transcription factors were unrelated to quiescent state switch.

Previous studies on other types of stem cells shown that the differentiation accompany cell fate decisions can occur in the lengthening G1 phase [21, 22]. But cells in G1 phase of the cell cycle are more prone to differentiate compared with S or G2/M phases [23]. There raises two questions: Why do progenitors prefer to make cell fate decisions in the G1 phase rather than in other cell cycle phases? One possibility is that the G1 phase of cell cycle has lower activation of molecular signals compared to the S and G2/M phases. Since cell fate decisions involve multiple lineages, simply signals of one process may avoid the interaction effects that can lead to mistakes in complex fate decision process. Further, this can ensure the mitosis is conducted correctly. Additionally, cells in the quiescent state have no specific tasks, so the activated state of the G1 phase serves as a cue for the cells to make a choice between fate decisions or self-renewal, facilitating the initial change in cell states. Furthermore, the S and G2/M phases provide a checkpoint of conversion that prevents incorrect fate decisions due to unreliable stimulation. Manipulating cell-cycle states could potentially enable control over differentiated cells. Secondly, why dose lineages commitment of differentiation in HPCs precedes cell division of proliferation? Mistake in proliferation may cause the generation of numerous abnormal cells, whereas a mistake in lineage commitment only affects a single cell. Lineages commitment was strictly orchestrated by extrinsic signals (cytokines) and intrinsic regulators (fate decision genes), that reduced the likelihood of errors. The disadvantage of an abnormal original cell was much smaller than that of a differentiated and committed cell. Committed progenitors can directly proliferate with high efficiency, satisfying the physiological requirements. In summary, the precise sequential control of differentiation and proliferation efficiently facilitate the urgent and stable requirements of specific cells during hematopoiesis.

Neutrophils are the primary and rapid responders during infections, capable of quickly generating an enormous number of mature neutrophils for immune defense [24, 25]. Our study showed that the continuous differentiation process of neutrophils is the shortest in the hematopoietic hierarchy. The progenitors quickly complete differentiation and proceed to proliferation in the second stages. CSF3R, the receptor of G-CSF, mobilizes progenitors from the bone marrow to peripheral blood. When G-CSF activates CSF3R, progenitors transition from a quiescent to an activated state, initiating the process of self-renewal or lineages commitment. Our study revealed that the expression characterization of CSF3R indicated that the initial mobilization of MPCs can quickly differentiate into neutrophils with minimal signaling changes. These characteristics of differentiation, the shortest differentiation process and minimal signaling changes, enable neutrophils to act as primary and rapid responders against infections, quickly generating an enormous number of mature neutrophils. The ’ultra-short-pathway with minimal-signaling’ strategy maximized bioenergetic and temporal efficiency by reducing both differentiation steps and regulatory molecule requirements. This mechanistic paradigm provides a molecular basis for understanding the innate immune system’s rapid-response capability. A two step pipeline for neutrophil generation is faster and more efficient than a lengthy procedure. The ’one-step’ commitment mechanism of NMPs (NMPs) challenges the conventional dogma that hematopoietic lineage commitment must undergo multiple transitional stages. This discovery demonstrated that during hematopoiesis, progenitor cells can adopt distinct commitment strategies to achieve both higher efficiency and functional adaptability—particularly evident in neutrophils’ rapid stress response during infection. Our study demonstrated that **lineage-specific efficiency stratification** existed during lineage commitment, suggesting the **paradigm shifts from a ’unified hierarchy’ to a ’functional-adaptive topological architecture’ model** during hematopoiesis.

The abnormal state and proliferation of HSCs are widely recognized as the key factors in hematologic malignancies [26]. However, the key genes or mechanism underlying some subtypes remain unclear. The main challenge and limitation lie in identifying aberrant genes or molecular signaling, as multiple activation states occur during hematopoiesis, including self-renewal, lineages commitment, proliferation. The complexity of these dynamic biological events makes it difficult to pinpoint the specific disease-associated genes and signaling changes. Untill now, the chronological order and states of these cellular activation events have been uncertain. In this study, we achieved a significant breakthrough in understanding cell cycle kinetics during hematopoiesis. By clearly distinguishing these dynamic biological events, we revealed the dynamic changes in cell cycle states and regulated genes. Our previously results clear identified the fate-decision genes of lineage commitment. By combining these two efficienct datasets and models, we can easily identify the aberrant genes or molecular signaling by comparing healthy and diseased individuals. The success of overcoming hematologic malignancies is now within reach.

Overall, we revealed the cell cycle kinetics of progenitor lineages and identified a specific factor, CDCA7, which is critical for DNA replication initiation during hematopoiesis. CDCA7 would be an effective target for drug development and leukemia therapy through controlling cell cycle activity. Our study further confirmed the shortest differentiation trajectory of neutrophils, facilitate their role as rapid responders during infections. This conclusion demonstrated that efficiency optimization in HPCs commitment can be achieved through pathway simplification rather than increased regulatory complexity. We provide detailed insights into the cell cycle kinetics of HPCs, revealing the chronological order of their transformation from differentiation to proliferation. Although multipotent self-renewal of HPCs does not occur after lineages commitment is completed in adult peripheral blood, proliferation does. Collectively, the cell cycle kinetics of hematopoiesis perfectly explains the physiological phenomenon of immunity. This study repensents a significant breakthrough in the fundamental discipline of immunity development. Often, the truth is simple, yet we have been confused for a long time.

## Methods

### HPC enrichment and ScRNA sequencing

Original peripheral blood HPCs from quiescent donors who had not received any treatment (11 individuals) and mobilized (G-CSF) donors (three individuals) were enriched using a surface marker panel (Lineage−) via negative deletion [10].10× Genomics RNA sequencing was performed on human peripheral blood enriched HPCs. In total, 27,566 CD34+ progenitor cells were extracted and re-analyzed to evaluate diversity, resulting in 17 clusters, including 21,244 mobilized and 6,322 quiescent progenitors.

### ScRNA sequencing data processing

Single-cell unsupervised dimensionality reduction and clustering functions were performed using Seurat [27]. Progenitor definition and annotation were followed previous results [10]. Gene Ontology (GO) enrichment and Kyoto Encyclopedia of Genes and Genomes (KEGG) analysis of genes were conducted using the clusterProfiler package in R software[28]. Score of cell cycle phases were analyzed by Seurat.

## Authors’ contributions

Y.Y. conceived the original idea, designed the study, and conducted the bioinformatics and statistical analyses of the sequencing data. Y.Y. wrote and revised the manuscript. All authors approved the final draft of the manuscript.

## Declaration of interests

The authors declare no competing interests.

## Notes

### Competing Interest Statement

The authors have declared no competing interest.

### Summary of Updates

1, Results were updated. 2, Graphical Abstract was added; 3, Disscussion was revised.

